# Low Intestinal Doses of Botulinum Neurotoxins types A and B favour infection by *Salmonella* and *Shigella* without the flaccid paralysis of botulism

**DOI:** 10.1101/2024.10.26.620416

**Authors:** Federico Fabris, Paola Brun, Aram Megighian, G Bernabè, Ignazio Castagliuolo, Ilenia Drigo, Luca Bano, Florigio Lista, Maria Lina Bernardini, Cesare Montecucco, Ornella Rossetto

## Abstract

Botulism is a life-threatening disease characterized by a descending flaccid paralysis caused by a protein neurotoxin (BoNT) released by different anaerobic bacterial species of the genus *Clostridium*. The paralysis results from blockade of neurotransmitter release from the terminals of peripheral cholinergic, skeletal and autonomic neurons exerted by BoNT through the cleavage of SNARE proteins, which are essential for neuroexocytosis. Here, we investigated the effect of different doses of BoNT serotypes A and B, the serotypes most commonly associated with human botulism, on enteric nervous system neurons which play an important role in gut health and physiology. We found that BoNT/A and BoNT/B enter cholinergic neurons where they cleave SNARE proteins even at doses that do not cause signs of flaccid neuroparalysis. However, these low BoNT doses favour the invasion and infection of the mouse body by *Salmonella thyphimurium* and *Shigella flexneri*. This may have significant animal health implications.

## Introduction

Botulism is a neurointoxication disease characterized by a peripheral descending flaccid paralysis with respiratory deficit and autonomic dysfunctions that can be life-threatening^1–4^. In fact, botulism is caused by botulinum neurotoxins (BoNTs), released by different species of bacteria of the genus *Clostridium*, which block neurotransmitter release from neurons for weeks to months depending on the type of BoNT.

BoNTs are 150 kDa proteins consisting of a 50 kDa light chain (L) and a 100 kDa heavy chain (H) linked by a single interchain disulfide bond. HC consist of a N-terminal translocation domain (HN, 50 kDa) and a C-terminal receptor-binding domain (HC, 50 kDa). BoNTs target cholinergic presynaptic neuron terminals via HC binding to polysialogangliosides and to specific presynaptic membrane proteins that mediate their endocytosis inside synaptic vesicles. These vesicles are used by the BoNTs as Trojan horses to reach the cytosol where the LC cleave at single peptide bonds one or more of the three SNARE proteins: VAMP (vesicle associated membrane protein), SNAP-25 (Synaptosome Associated Protein-25) and syntaxin. Their cleavage impairs the release of the content of synaptic vesicles and granules ^5–8^. This biochemical lesion is reversible with time, owing to degradation of the LC, and does not cause neuronal death, making the flaccid paralysis of botulism reversible. Mechanically ventilated patients will recover within a period dependent on the dose and type of BoNT^1,9^.

In fact, BoNTs are produced in eight different serotypes (BoNT/A-G and BoNT/X) and a large number of subtypes that exhibit a large range of toxicity in vertebrates^6,7,10–13^. In addition, an increasing number of BoNT-like proteins that target insects or yet non-defined animal targets have been discovered^14–16^.

BoNT/A, /B, /E, and rarely F, are the toxin serotypes causing botulism in humans whereas BoNT/C and BoNT/D and their mosaic forms C/D, D/C affect animals^1–4, 17,18^. Foodborne and infant botulism are the most common form of the disease in humans and they develop following ingestion of food containing pre-formed active toxin in complex with accessory proteins or neurotoxic Clostridia spores, respectively. The botulinum complex, known as progenitor toxin complex (PTC) is released by the bacterium and consist of a BoNT molecule interlocked to a non-toxin non-haemagglutinin (NTNH) component and to several nontoxic neurotoxin-associated proteins (NAPs) of various sizes^19^. The PTC is much more resistant than the free BoNT to the harsh conditions present in the gastrointestinal tract (GI), and its oral median lethal dose is several fold lower than that of free BoNT^12,20^. Once in the small intestine the PTC dissociates and the BoNT crosses the enteric epithelial barrier in a yet poorly understood way. From the intestinal mucosa the BoNT is believed to diffuse locally reaching the lymphatic and blood circulations which deliver the toxin to cholinergic nerve terminals of the body blocking acetylcholine (ACh) release and causing the flaccid paralysis characteristic of botulism^21,22^.

In humans, this paralysis begins with ptosis, diplopia and dysphagia, and in a descending mode, it diffuses reaching all skeletal muscles, including those involved in respiration^1,3,9,23,24^. Also neurotransmission of autonomic nerve terminals is impaired by BoNTs, particularly by type B, which cause autonomic dysfunctions including alterations of the heart rhythm, orthostatic and supine hypertension with impaired baroreflex function, dry mouth and eyes, hypo-/anhidrosis, flaccid bladder, urinary retention and erectile dysfunction. In addition, constipation and diarrhea are present in most cases and may progress to adynamic ileus^25–27^. Constipation is likely to be due to impaired neurotransmission of enteric neurons governing contraction of the circular and longitudinal smooth muscles that control peristalsis.

Although intestinal dysfunction is a major symptom of botulism, until now the small intestine has been considered only as the transient route of entry of BoNTs into the body. Little attention has been paid to the local effect on the Enteric Nervous system (ENS), despite the fact that ENS together with the extrinsic vagal and spinal nerves, orchestrate many vital functions and the fact that the concentration of BoNT is higher in the intestine in botulism, and is diluted after crossing the intestinal barrier. The ENS is characterized by a highly developed array of neurons and glia organized in two distinct ganglionated plexuses and their interconnecting neural pathway^28^. The neurons of the myenteric (or Auerbach) plexus coordinate smooth muscle contractions and the neurons of the inner submucosal (or Meissner) plexus control gut secretions, nutrient absorption, and local blood flow^29^. A recently acquired major knowledge is that many enteric neurons interact directly with local immune cells including enteric macrophages, innate lymphoid cell, dendritic cells, glial cells and several types of lympocythes. They form neuro-immune cell units that govern intestinal immune response, and consequently the defence against the invasion of intestinal pathogens ^30–33^. Moreover enteric cholinergic neurons stimulate goblet cells to produce and release apically mucins and other proteins forming the mucus layers that cover the luminal side of the GI tract^28^ and is a first barrier to prevent bacterial invasion of the intestinal mucosa.

About 80% of enteric neurons of the ENS are cholinergic and are, therefore, a natural first targets for BoNT intoxication^29,33^. Among their known functions, enteric cholinergic neurons control intestinal motility, mucus secretion from the goblet cells and, interact with immune cells thus modulating the defence against intestinal pathogens^34–37^.

Here, we have used an *in vivo* mice model of food-borne botulism type A and B (the two forms that account for most of the fatal cases of human botulism) and have investigated the ability of orally administered BoNT/A or BoNT/B to intoxicate enteric neurons. In order to simulate the *in vivo* situation, we used culture supernatants of *C. botulinum* type A and B (sBoNT/A and sBoNT/B) administered by oral gavage in mice. We analyzed their local enteric and systemic effects using different BoNT doses, thus mimicking what happens in human BoNT poisoning. We have found that low doses of sBoNTs that do not cause any overt symptom on respiratory and skeletal neurons, do instead alter intestinal functions and allow the tissue invasion by important intestinal microbial pathogens such as *Salmonella typhimurium* and *Shigella flexneri*. These results highlight ENS as the first target in food-borne and infant botulism and suggest the medically relevant possibility that even minor doses of BoNT are pathologically important as they alter intestinal neuronal functions and facilitate the entry of intestinal microbes that may disseminate and cause infectious diseases.

## Results

### Effect of BoNT/A and BoNT/B culture supernatants on primary cultured neurons

In order to study the effects of BoNT/A and BoNT/B on the small intestine of mice under conditions simulating food-borne and infant botulism, *C. botulinum* supernatants of cultures of strains ATCC19397 and ATCC27765 (sBoNT/A and sBoNT/B, respectively) (fig. 1a) were delivered to mice by oral gavage. Before mice delivery, sBoNT/A or sBoNT/B were characterized for their content of active toxin type A and type B, respectively, by comparing their proteolytic activity with those of isolated BoNTs of known concentrations in cerebellar granule neurons (fig. 1b-c). Both sBoNTs showed proteolytic activity, which was detected by the ratio of whole SNAP25 (e-SNAP25) to cleaved SNAP25 (cl-SNAP25) (fig. 1b) for sBoNT/A, and by the ratio of whole VAMP2 (e-VAMP2) to cleaved VAMP2 (cl-VAMP2) for sBoNT/B (fig. 1c). This comparison and the determination of the 50% dose causing mouse lethality (mLD_50_) in the classical mouse bioassay led to the identification of the doses of sBoNT/A and sBoNT/B to be used (fig. 1a).

**Figure 1.**
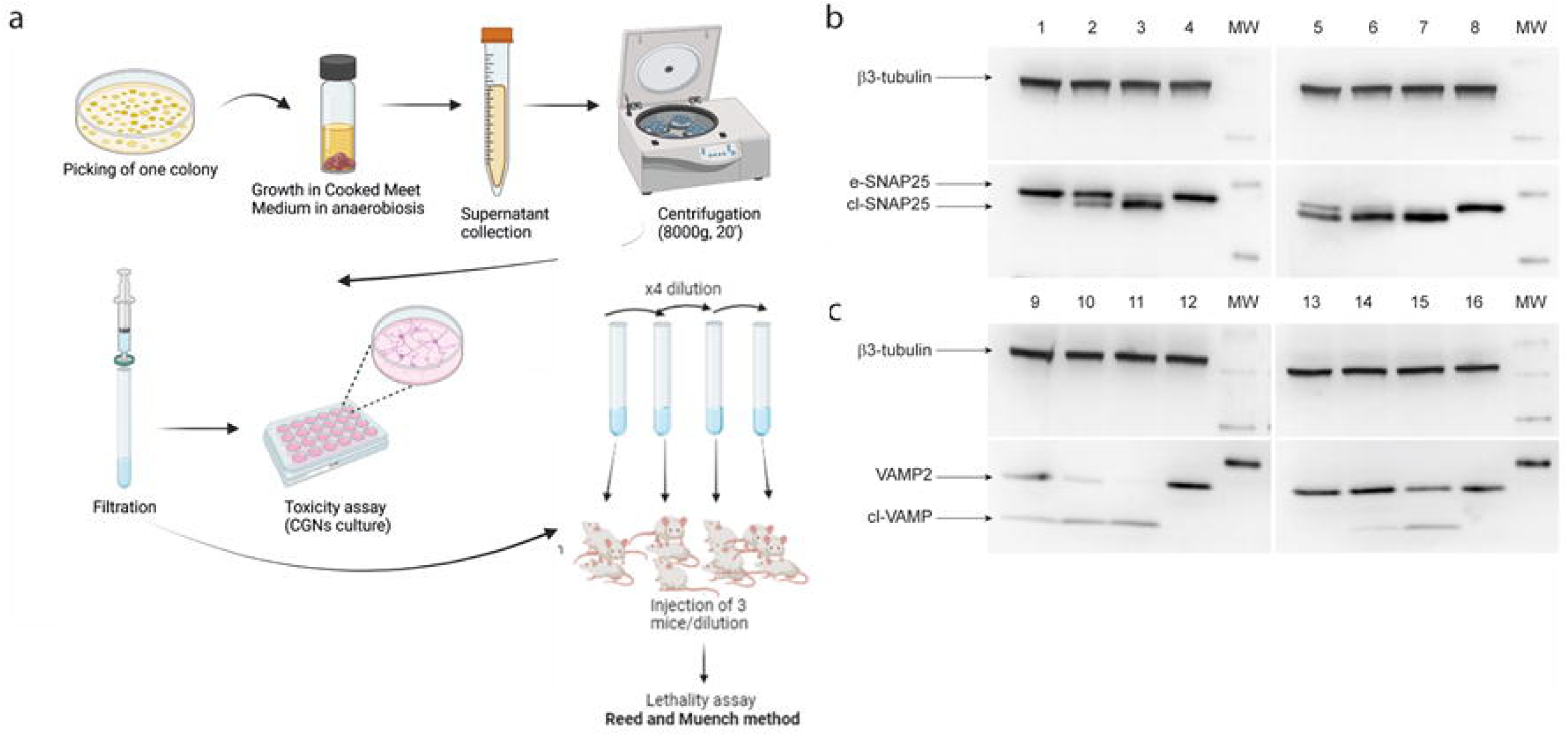
Protocol for culture and extraction of BoNT complexes and determination of their toxicity. **a,** schematic representation of the culture protocol for *C. botulinum* to obtain supernatants containing BoNT complexes (sBoNT/A, sBoNT/B). Once obtained, the supernatants were tested in vivo and in vitro to assess their toxicity. **b,** Western blot on CGNs lysates to evaluate the toxicity of sBoNT/A. The comparison was made between CGNs treated with different concentrations of purified BoNT/A (left column, upper panel; 1-3: 0.1 pM, 1 pM, 10 pM, respectively; 4: vehicle) or different volumes of sBoNT/A (right column, upper panel; 5-7: 0.1 µL, 1 µL, 10 µL, respectively; 8: vehicle). **c,** Western blot on CGNs lysates to evaluate the toxicity of sBoNT/B. The comparison was made between CGNs treated with different concentrations of purified BoNT/B (left column, lower panel; 9-11: 0.05 nM, 0.5 nM, 5 nM, respectively; 12: vehicle) or different volumes of sBoNT/B (right column, lower panel; 13-15: 0.05 µL, 0.5 µL, 5 µL, respectively; 16: vehicle). The cartoon in **a** was created using Biorender.com.

### Systemic effects of ingested BoNT/A and BoNT/B culture supernatants

Oral gavage of different volumes of sBoNT/A or sBoNT/B was performed and we used a “high” dose that determined clinical botulism (ruffled fur, hind limb paralysis, laboured breathing and death) and a “low” dose that did not elicit any of the manifestations of the flaccid paralysis of botulism. Qualitative external observations were accompanied by quantitative measurements of the main motor function parameters involved in botulism (fig. 2a-d). Compound Muscle Action Potential (CMAP) electromyography was used to assess the neurotransmission at the neuromuscular junctions (NMJ) in the whisker pad (fig. 2a) and in the gastrocnemius muscle (fig. 2b). The peak amplitude of CMAP was markedly reduced in both muscles three days after oral gavage of animals with the high dose of sBoNT/A or sBoNT/B with respect to controls treated with vehicle, consistent with the flaccid paralysis of botulism. At variance, the CMAP peak amplitude value was unchanged at both three and seven days after oral treatment of mice with the low dose of sBoNT/A or sBoNT/B (fig. 2a-b). Similar results were obtained with the Rotarod test (fig. 2c), which showed a significant reduction in the performance of high-dose treated mice, in comparison with those receiving the low-dose treatment.

**Figure 2.**
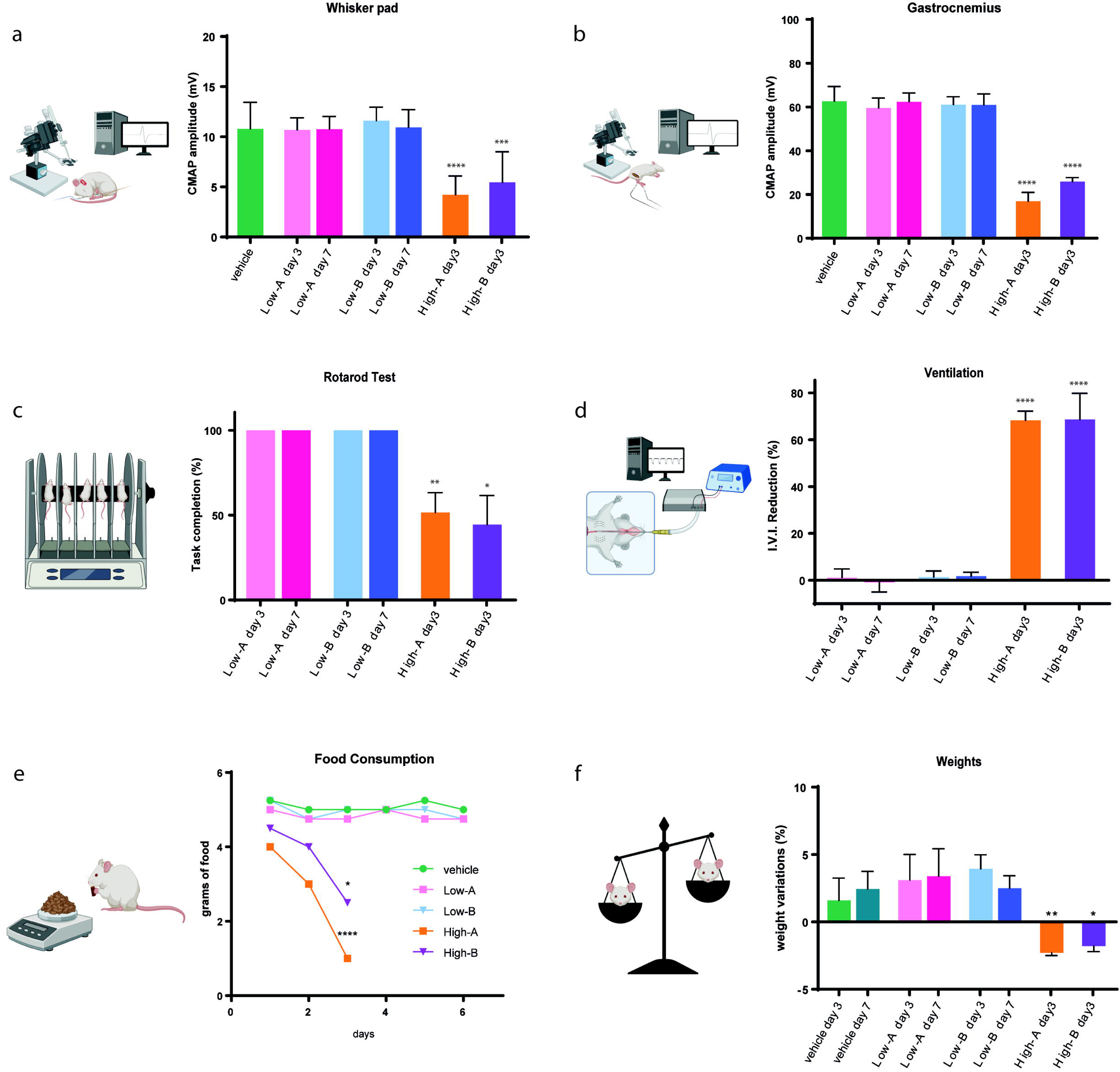
Oral intoxication with BoNT causes impairments in the well-being of mice treated with high doses of toxin, but not in those treated with low doses. **a.** CMAP analysis in the whisker pad. CMAP recording in the whisker pad is performed by stimulating the facial nerve (see cartoon on the left). The data come from mice treated with vehicle only (in green), mice intoxicated with a low dose of sBoNT/A (in pink at three days, fuchsia at seven days) or sBoNT/B (in light blue at three days, blue at seven days), and mice treated with high doses of toxins (sBoNT/A in orange, sBoNT/B in purple, both at three days). The graph shows CMAP amplitudes (in mV). **b.** CMAP analysis in the gastrocnemius muscles. CMAP recording is performed by stimulating the sciatic nerve (see cartoon on the left). Similarly, the data come from mice treated with vehicle (green), intoxicated with a low dose of sBoNT/A (pink, fuchsia) or sBoNT/B (light blue, blue), or mice intoxicated with high doses of toxins (sBoNT/A in orange, sBoNT/B in purple). The graph shows CMAP amplitudes (in mV). **c.** Motor activity evaluation via Rotarod test. The Rotarod test evaluation (cartoon on the left) is performed using a standard protocol (see materials and methods). Results are reported as the % of the test completed by mice treated with low doses of sBoNT/A or /B (pink and fuchsia for the former, light blue and blue for the latter) or high doses of toxins (in orange and purple, respectively). **d.** Ventilation analysis. The graph shows the percentage variation in ventilation capacity observed in mice treated with low doses (pink and fuchsia for sBoNT/A, light blue and blue for sBoNT/B) or high doses (orange for sBoNT/A, purple for sBoNT/B) of toxin, compared with the results from the same mice pre-treatment. Ventilation was measured exploiting pressure variations in mice lower esophagus using a gavage needle connected to a plastic tube (cartoon on the left). **e.** Measurement of food intake in BoNT-intoxicated mice. The graph shows fluctuations in food consumption by mice treated with vehicle only (green), low doses of sBoNT/A or /B (pink and light blue, respectively), or high doses of the same toxins (orange and purple, respectively). **f.** Weight variation analysis. The data are reported as weight changes in animals treated with vehicle only (light green, dark green), low doses of sBoNT/A (pink at three days, fuchsia at seven days), sBoNT/B (light blue at three days, blue at seven days), or high doses of toxins (at three days, sBoNT/A in orange, sBoNT/B in purple). Statistical analysis was done by using One-way analysis of variance (ANOVA) with Tukey’s post-hoc test and multiple comparisons for graphs a, b, e and f, while Wilcoxon signed-rank test was applied to data of graphs c and d, with 100 and 0 as expected values, respectively. Data were considered statistically different when *p < 0.05, ** p < 0.01, *** p < 0.001, **** p < 0.0001. All cartoons were created using Biorender.com.

A major symptom of botulism is the impairment of respiration due to the systemic BoNT diffusion that reaches and inhibits neurotransmission of the NMJ of respiratory muscles. Lung ventilation in the different groups of mice was significantly reduced three days after the treatment with high dose of sBoNT/A or sBoNT/B, whereas this parameter remained unchanged at three and at seven days after oral gavage of mice with low dose of neurotoxins supernatants (fig. 2d).

These functional assays were paralleled by imaging of the NMJ of the whisker pad muscles, diaphragms and gastrocnemi staining for cl-SNAP25 sBoNT/A-treated animals or and with an antibody that specifically recognizes cleaved VAMP-1/2 in the NMJs of sBoNT/B-treated animals used for the experiments reported in fig.1. As shown in fig. S1, high percentage of NMJs showing either cl-SNAP25 or cl-VAMP signal was evident only in animals treated with high dose of supernatants, whereas significant less proteolytic activity was observed in muscles from animals treated with low dose of supernatants. In addition, fig. S2 shows that no active toxin proteolytic activity was detected in the serum of animals treated with the low dose of sBoNT/A or sBoNT/B using the well-established Endopep-MS method^38–40^. These results suggest that the amount of active toxin present in the general body circulation of mice treated with the low dose of supernatants is not sufficient to affect neurotransmitter release from motor neuron axon terminals.

### Low Dose sBoNT/A or sBoNT/B cleave SNAREs within enteric cholinergic neurons impairing gut mobility

Despite the absence of signs of flaccid peripheral paralysis in mice treated with a low dose of sBoNT/A and /B, we tested the possibility that SNARE cleaved by these neurotoxins within enteric neurons could affect gut physiology. Two of the most reported enteric symptoms of foodborne botulism are constipation and diarrhea, so we measured the faeces transit time and faecal moisture content in mice gavaged with sBoNT/A or sBoNT/B. By using the charcoal motility test, we observed a significant slowdown in faeces transit time after the treatment with low dose of sBoNT, inferring an impairment of intestinal peristalsis, as early as day three that persisted seven days after the treatment (fig. 3a). These data were accompanied by a significant increase in fecal pellet humidity, which was observed both in low-dose and high-dose treated mice (fig. 3b). As cholinergic neurons of the myenteric plexus are known to control the intestinal motility via release of acetylcholine (ACh) that activates contraction in the longitudinal and circular muscles^36,41^, we checked for BoNTs proteolytic activity in the Enteric Nervous System (ENS). The myenteric plexus is extensively interconnected with the submucosal plexus located within the connective tissue of the submucosa, which controls secretory functions, absorption and blood flow. Therefore, both myenteric and submucosa plexus were isolated by peeling the different layers of the small intestine of intoxicated animals. As the initial part of the upper small intestine is indicated as the primary site of adsorption of BoNTs^21,42^ we focused on this portion of the intestine. The intracellular metalloproteolytic activity of the toxin was detected as previously described^43^. The panels c and d of fig. 3 show that sBoNT/A and sBoNT/B cleave SNAP-25 and VAMP, respectively, in both the myenteric and submucosal plexuses. Their proteolytic activity was mainly detected in cholinergic neurons, as indicated by the co-localization of cleavage signals with the cholinergic marker vesicular acetylcholine transporter (VAChT). Co-localization with cleaved SNAREs was extensively observed also in neurons containing Substance P (SP), whilst very little or no signal of SNARE cleavage was detected in tyrosine hydroxylase (TH) or vasoactive intestinal peptide (VIP) containing neurons (Fig. S3). These data are in agreement with the selectivity of BoNTs for cholinergic neurons. Indeed, most of myenteric excitatory motor neurons that release ACh as primary neurotransmitter, contained in synaptic vesicles, also release other molecules, such as SP contained in electron dense presynaptic granules (fig. S3). Therefore, BoNT/A and BoNT/B appear to be rather selective for ENS cholinergic neurons when the intoxication occurs with low doses of toxin.

**Figure 3.**
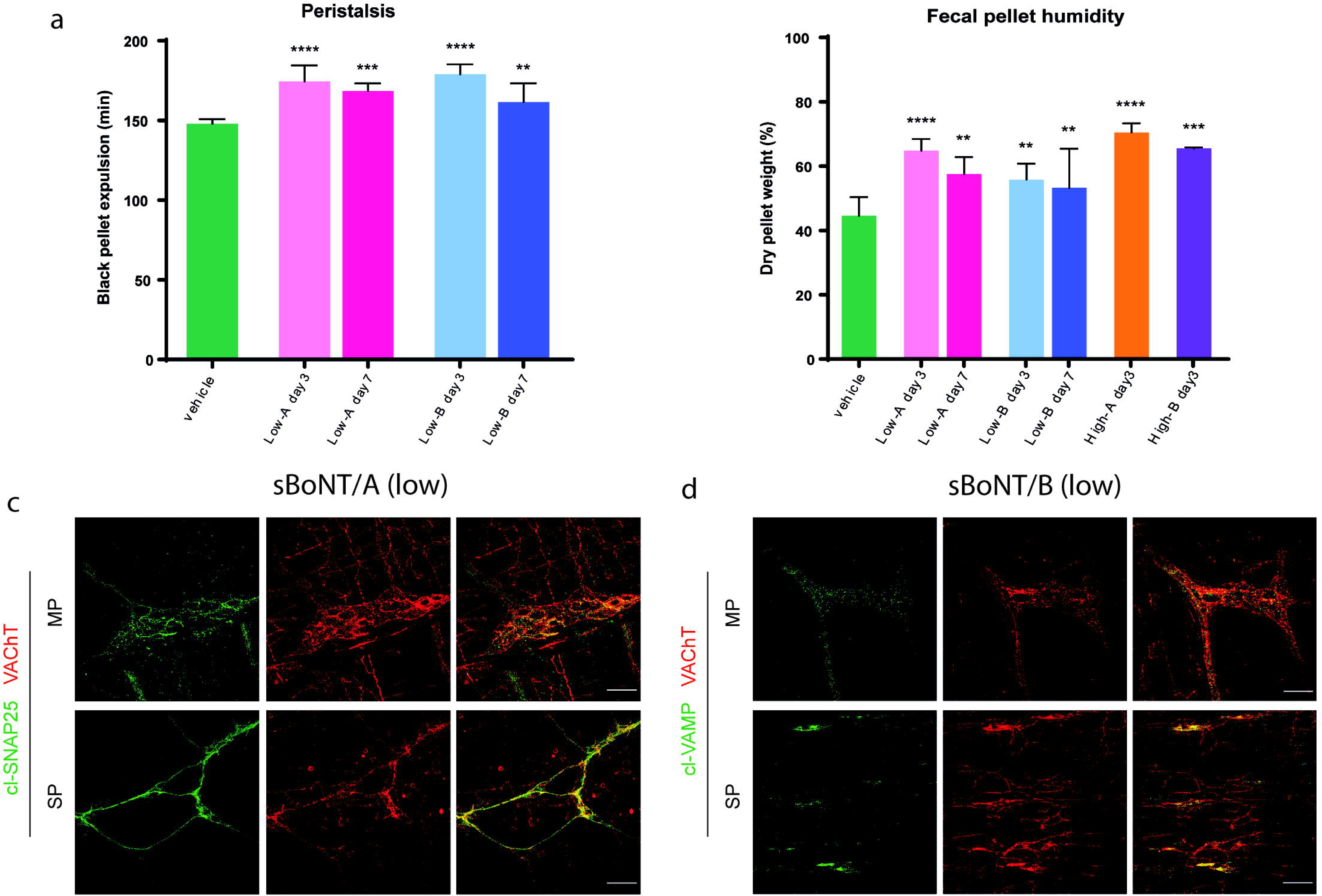
BoNTs proteolyze their SNARE targets, causing impairments in gut physiology. **a.** Measurement of peristalsis speed using the charcoal motility test. The data show the time measured (in minutes) from the moment of gavage with a charcoal and gum arabic bolus to the expulsion of the first black fecal pellet. The data refer to mice treated with low doses of sBoNT/A (in pink at three days, fuchsia at seven days) or sBoNT/B (in light blue and blue), or one week before intoxication (referred to as vehicle, in green). **b.** Measurement of fecal pellet moisture. The data show the moisture content (reported as percentage of faecal weight loss) in the feces of mice treated with low doses of toxin (sBoNT/A in pink and fuchsia; sBoNT/B in light blue and blue) or high doses of toxin (sBoNT/A in orange; sBoNT/B in purple). The moisture content in the fecal pellets of the same mice one week before intoxication is reported in green (vehicle). **c.** cl-SNAP25 staining in enteric ganglia. cl-SNAP25 staining (in green) is found in both the myenteric plexus (upper panel) and submucosal plexus (lower panel) of mice intoxicated with low doses of sBoNT/A. High colocalization (in yellow) with cholinergic neurons (stained for VAChT, in red) is observed. **d.** cl-VAMP staining in enteric ganglia. cl-VAMP staining (in green) is found in both the myenteric plexus (upper panel) and submucosal plexus (lower panel) of mice intoxicated with low doses of sBoNT/B. High colocalization (in yellow) with cholinergic neurons (stained for VAChT, in red) is also observed in this case. Immunofluorescence images were acquired at 20X magnification. Scale bar: 50µm. Statistical analysis was performed using one-way analysis of variance (ANOVA) with Tukey’s post-hoc test and multiple comparisons. Data were considered statistically significant when *p < 0.05, ** p < 0.01, *** p < 0.001, **** p < 0.0001.

Taken together, these results clearly indicate that BoNT/A and BoNT/B are capable of intoxicating enteric cholinergic neurons of both myenteric and submucosal plexuses, leading to at least two alterations of intestinal physiology at doses that do not cause the botulism flaccid paralysis. This implies that the ENS is the primary site of action of botulinum neurotoxins types A and B during foodborne and infant botulism and highlight that a low dose of BoNT can be a potential cause of intestinal dysfunction even in the absence of botulism symptoms.

### sBoNT/A AND sBoNT/B favour bacterial translocation across the intestinal barrier

Enteric cholinergic neurons interact with immune cells forming neuro-immune units that control the protective response against intestinal pathogens^30,44,45^. Given that the blockade of enteric neurotransmission by BoNT/A and BoNT/B cause constipation and fluid faeces, we hypothesized that the blockade of cholinergic neurotransmission also impair immune host defence thus increasing enteric bacteria translocation with ensuing dissemination in the body. This hypothesis was tested using two prototypical enteric pathogens. *Salmonella enterica serovar typhimurium* is able of paracellular translocation through the intestinal barrier whilst *Shigella flexneri* performs intracellular infection and transcellular dissemination^46,47^. As shown in panel A of fig. 4, mice pre-treated with toxins or vehicle were administered with suspensions of *S. typhimurium* (1 × 10^8^ CFU) or *S. flexneri* (1 × 10^7^ CFU)^48,49^ by oral gavage. Five days after the bacterial inocula, mice were euthanized and the Peyer’s patches (PP), liver and spleen were removed and homogenized. Tissue homogenates were plated on bacteriological agar and bacterial CFU were enumerated to assess bacterial invasion and dissemination. As reported in fig. 4b, more *S*. *typhimurium* colonies were found in all the tissues obtained from sBoNT/A- and sBoNT/B treated mice as compared to the vehicle treated group denoting an increased entry of this pathogen in the mouse body. Similarly, in the Peyer’s patches collected from *S. flexneri* infected animals, we observed a higher number of bacteria CFUs in toxins treated mice than in the vehicle group (fig. 4b). No CFU of *S. flexneri* were found in the spleen and liver, independently from the pretreatment consistent with the different routes of infection of *Shigella* spp^49–51^ in comparison with *Salmonella* spp^48,52^. The invasion of these intestinal pathogens was also visualized by infecting mice with GFP-fluorescent bacteria. Fig. 4c shows that larger aggregates of bacteria were present in the PP of BoNT-treated mice than in vehicle mice. These results suggest that low concentrations of BoNT/A and /B, by impairing the neurotransmission of cholinergic enteric neurons, reduce the protective response of the host against the invasion of intestinal pathogenic bacteria.

**Figure 4.**
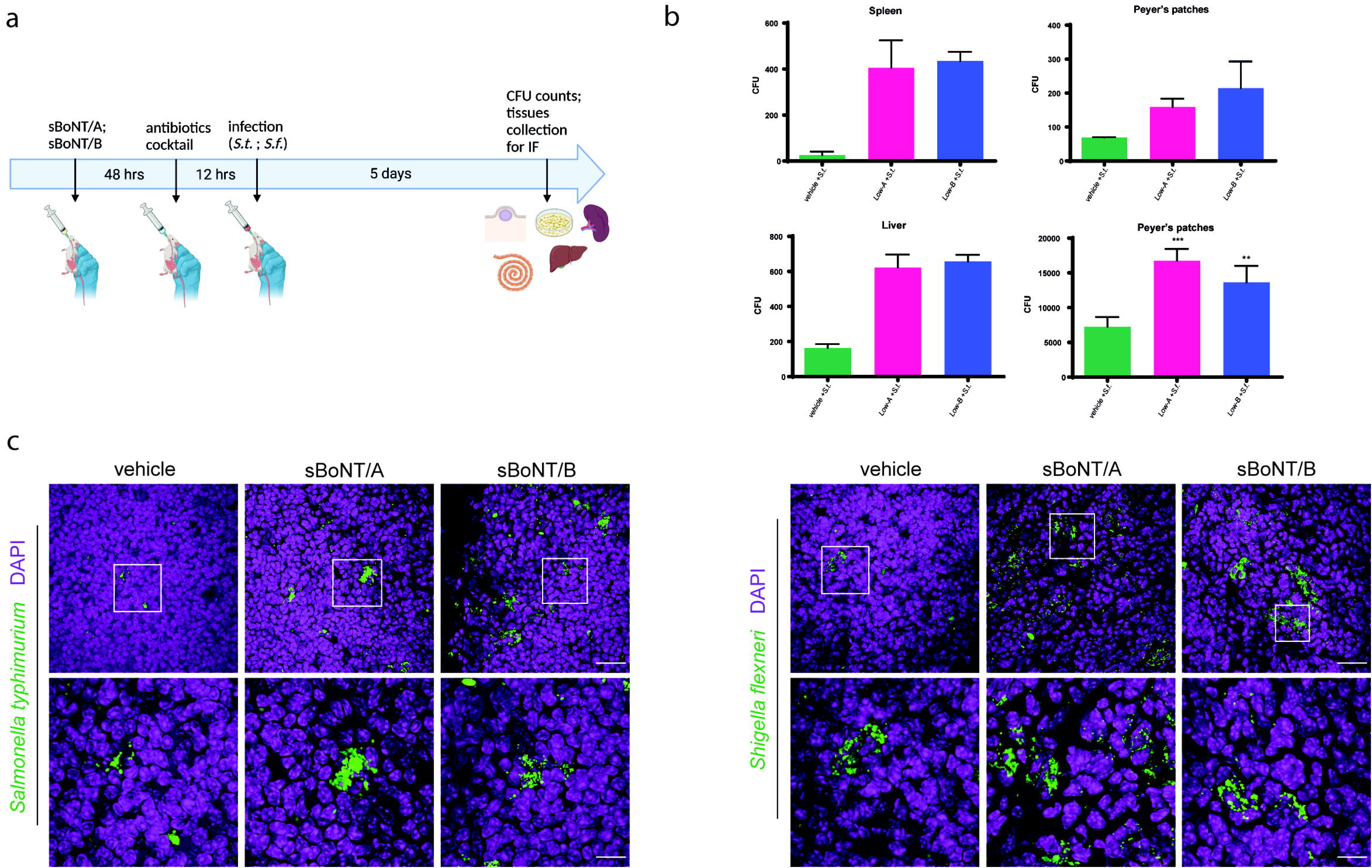
Evaluation of the invasiveness of foodborne pathogens in BoNT-intoxicated mice. **a.** Experiment workflow. 72 hours after intoxication with a low dose of sBoNT, mice are orally infected with *Salmonella typhimurium* (S.t.) or *Shigella flexneri* (S.f.). At the end of the experiment, five days later, tissues are collected for CFU counting and immunofluorescence experiments. **b.** CFU count of *Salmonella typhimurium* and *Shigella flexneri* in Peyer’s patches, spleen, and liver. For the *Salmonella typhimurium* infection model (top graphs, low-left graph), CFU counts for all three tissues are reported; for the *Shigella flexneri* infection model (low-right graph), CFU counts for Peyer’s patches are shown. CFU counts from mice pre-intoxicated with a low dose of sBoNT/A are shown in fuchsia, those from mice pre-intoxicated with a low dose of sBoNT/B are shown in blue, and mice infected with oral pathogens without pre-intoxication (vehicle) are shown in green. **c.** Staining of fluorescent bacteria in Peyer’s patches. Representative images of *Salmonella typhimurium*-GFP (left panel) and *Shigella flexneri*-GFP (right panel) in the Peyer’s patches of animals pre-treated with a low dose of sBoNT (middle and right images of each panel) or vehicle alone. Images were acquired at 40X (upper panels, scale bar: 25 µm) or 100X (lower panels, scale bar: 10µm). Statistical analysis was performed using one-way analysis of variance (ANOVA). Data were considered statistically significant when **p < 0.01, ***p < 0.001, ****p < 0.0001. The cartoon in **A** was created using Biorender.com.

The GI tract defence system operates via different lines of action. One of the most important line is the release of great mucus cargos by the goblet cells (GC) under the stimulus of ACh released by impinging neurons (fig. 5a). We analyzed this aspect in the intestine roll preparations obtained from mice infected with either *Salmonella* or *Shigella* with or without pre-intoxication with sBoNT. As shown in the enteric cryo-slices stained with the mucus marker WGA and the signal intensity quantification (fig. 5b-d), we observed that mice treated with sBoNT exhibited increased mucus retention along villi and in enteric crypts of the intestinal mucosa, compared to vehicle treated animals. Notably, the reduction of the enteric physical and chemical barrier leads to increased bacterial invasion^53,54^, as observed in our sBoNT-treated mice.

**Figure 5.**
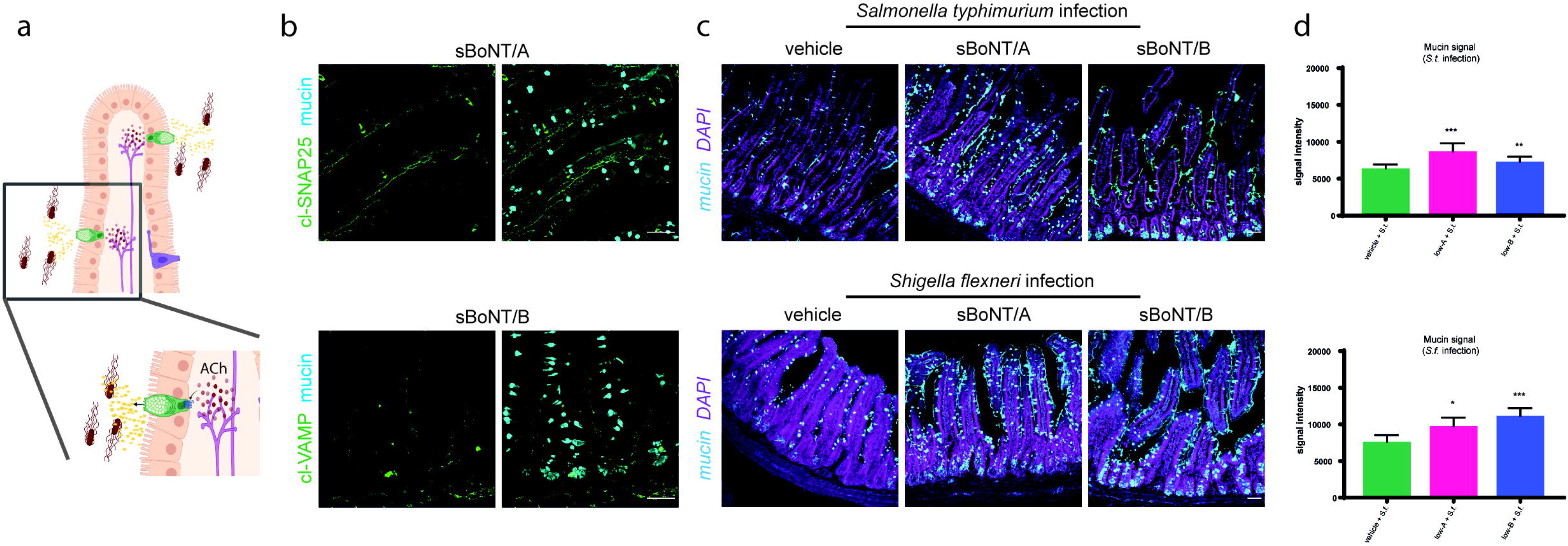
Evaluation of mucin secretion in response to foodborne infections in BoNT-intoxicated mice. **a.** Illustrative cartoon of cholinergically-stimulated mucus secretion in response to infections. **b.** Staining of cleaved SNAP25 (in green, upper panel) and VAMP (in green, lower panel) in axons near Goblet Cells (in blue) of mice post-intoxication with low doses of sBoNT/A (upper panel) or sBoNT/B (lower panel). Images were acquired at 40X magnification (scale bar: 25 μm). **c.** Staining of mucus within Goblet Cells of mice infected with foodborne pathogens. Mucus accumulation (shown by mucin staining, in blue) was evaluated in mice infected with *Salmonella typhimurium* (upper panel) or *Shigella flexneri* (lower panel), pre-intoxicated with low doses of sBoNT/A (middle images) or sBoNT/B (right images), or pre-treated with vehicle alone (left images). DAPI staining is shown in purple. Images were acquired at 20X magnification (scale bar: 25 μm). **d.** Quantification of intracellular mucus fluorescent signal. The graphs show the average fluorescence intensity within Goblet Cells of mice infected with *Salmonella typhimurium* (upper graphs) or *Shigella flexneri* (lower graphs). Data from mice pre-treated with vehicle are shown in green, while data from mice pre-intoxicated with low doses of sBoNT/A or sBoNT/B are shown in fuchsia and blue, respectively. Statistical analysis was performed using one-way analysis of variance (ANOVA) with Tukey’s post-hoc test and multiple comparisons. Data were considered statistically significant when *p < 0.05, **p < 0.01, ***p < 0.001. The cartoon in **a** was created using Biorender.com.

## Discussion

The main finding presented here is that a low dose of BoNTs, which is insufficient to cause any of the manifestations of peripheral flaccid paralysis, causes enteric changes that favour the possibility of an intestinal infectious disease. Currently, this event is most unlikely to be clinically reported as caused by BoNTs because poorly defined symptoms such as intestinal malaise and decreased intestinal motility are elicited that are more likely attributed to a variety of other causes^23,55–57^. However, this event is expected to be rather frequent particularly in countries where home prepared and canned food is a common practice. In addition, this can be deduced also by the description of literature reports of botulism outbreaks originated by large number of people sharing the same BoNT contaminated meal and resulting in various degrees of neuroparalysis from deadly to undetectable^55,56,58^.

Here, we document the possibility that low doses of BoNT cause local blockade of release of ACh through the specific cleavage of SNARE proteins within cholinergic enteric neurons. Given the documented role of cholinergic neurons in the defence of the organism against enteric invasion, their inhibition results in a reduction in some intestinal defensive systems. This is evidenced by an increase in intestinal permeability to bacteria, which eventually favours their translocation and causes diseases other than botulism, such as systemic inflammation with potential adverse consequences. The present work provides a proof of concept that botulinum neurotoxins at low doses, by intoxicating enteric neurons, allow dissemination of enteric pathogens. Here we have tested two paradigmatic bacteria, but it is likely that intoxicated enteric neurons fail in constraining enteric viruses, fungi and parasites^59^. Therefore, we posit that, although botulism is a rare disease, food poisoning with low doses of BoNTs, which remains undetected due to the lack of botulism symptoms, can be anything but rare and may cause infections with opportunistic bacteria.

Our study was performed by gavaging mice with low doses of BoNT/A and BoNT/B culture supernatants, to be as close as possible to what may happen in human cases of botulism. We have used BoNTs serotypes A and B because they are more frequently associated to human botulism but these effects may be observed with other serotypes as well, like BoNT/E and BoNT/F. The same consideration also applies to other BoNT serotypes with respect to animal botulism of other vertebrates: for example BoNT/C for birds particularly for those grown in intensive production facilities and BoNT/C and /D and their chimeras for bovines^18, 60–62^. Thus, the present study should be extended to test the effects of low doses of other serotypes on the infectivity of intestinal pathogens. Here, we intended to provide sufficient experimental evidence of a medically relevant possibility that has not been considered before to the best of our knowledge.

Our study is based on recent growing body of evidence indicating that the intestine wall is a site of extensive functional interactions between the nervous and the immune systems, as well as more generally of the intestinal defence. Indeed, these physiological interactions are essential for regulating the composition of intestinal microbiota and for preventing the invasion of bacteria, fungi and parasites from the intestinal lumen. Enteric neurons were recently shown to interact with various immune cells, like macrophages, innate lymphoid cells, dendritic cells and glial cells. This cooperation is central for shaping an intestinal homeostasis in the immune response to microbial pathogens present in the intestinal lumen^63–69^. In addition, the role of Ach released by cholinergic neurons on the stimulation of goblet cells to produce and release the component of the mucus layers has been extablished^54^.

Approximately 80 % of the neurons of the ENS are cholinergic^29^. Popoff and colleagues had previously found that the ENS can be labelled by high concentrations of fluorescent derivatives of the HC binding domain of BoNTs, which were shown to enter the mucosa via crossing the epithelial polarized monolayer of the intestine^70,71^. Based on this evidence, we postulated that BoNTs could enter and inactivate these neurons, thereby impairing enteric homeostasis. This hypothesis had become testable because we developed a method to visualize the cleavage of the SNARE inside neurons with a high sensitivity^43,73^. As expected, this procedure induced the flaccid peripheral paralysis of botulism when sBoNTs were administered at high concentrations, as documented by the quantification of a series of parameters that included CMAP in both the whisker pad and the gastrocnemius, Rotarod test and lung ventilation tests (fig. 2a-d). This approach enabled us to identify, by scaling down the doses of sBoNT/A and sBoNT/B, a dose affecting intestinal physiology without acting at distance (Figg. 2 and S1), most likely due to the toxin dilution associated with diffusion in the body. Indeed, in serum of low-dose intoxicated mice BoNTs were not detected using the highly sensitive endopep-MS assay and by the low levels of cleaved SNAREs in peripheral skeletal muscles, which was below the concentration required to block neuroexocytosis (fig. S2 and S1). SNAP-25 and VAMP cleavage by BoNT/A and /B, respectively, was detected within cholinergic neurons of both the myenteric and the submucosal plexus of the upper small intestine (fig. 3). This is the first time that BoNT/A and BoNT/B are shown to act directly in enteric neurons *in vivo* and, more relevant, at a low subclinical dose. Previous studies have used tissues or ligated intestines and focused on enteric neuron binding by fluorescent BoNTs or their fluorescent binding domains, which require great amounts of protein to be detectable ^22,70–72^. Our experimental approach was based on the use of polyclonal antibodies able to recognize with high sensitivity the cleavage product of BoNT/A and /B. The immunofluorescence signal is amplified by the enzymatic activity of the toxin, making these signals a smooking-gun proof of BoNTs impairment of ENS neurotransmission^43,73^. Differently from previous BoNT binding studies, in our experimental conditions non-cholinergic enteric neurons (cathecolamine- or VIP-immunoreactive neurons) were not targeted by BoNT/A and BoNT/B probably due to the low toxin concentration (fig. S3), which indicates a high toxin affinity for cholinergic neurons in comparison with other neurons in the ENS, a condition observed also in the PNS. These results define the gut not as a simple gateway for botulinum toxin entry but as their first site of action in foodborne and infant forms of botulism.

Taken together the present findings call for a reconsideration of botulinum neurotoxins, in the sense that they should also be viewed as *indirect enteric immunotoxins*, acting in minute amounts to depress the activity of enteric mucosal immune cells. Such an inhibitory effect would confer an evolutionary advantage to the Clostridia, as it would create the conditions for invasion of the body by intestinal pathogens, which would subsequently decrease the fitness of the intoxicated/infected animal in the natural environment. The death of an animal creates an anaerobiotic cadaver where anaerobic Clostridia can proliferate rapidly before the exhaustion of resources, subsequently transforming into spores and diffusing into the environment over long distances. In this sense, BoNTs would join the considerable number of bacterial toxins that impair the immune system in one way or another. Finally, these novel findings suggest that BoNTs can be employed as tools to investigate the neuro-immune cross-talk in mucosal tissues. Exciting opportunities in which the ENS could be used as a therapeutic target for common GI diseases could also be considered, as the further elucidation of such mechanisms could facilitate more therapy-related advances, ultimately changing the treatment approach.

## Material and Methods

### Ethical statement

All procedures were performed under general anesthesia and analgesia, when required. Tissue sampling was carried out with animals sacrificed by transcardial perfusion under deep anesthesia, or by cervical dislocation. All the experiments that involved animals were carried out in accordance with National laws and policies (D.L. n. 26, March 14, 2014), with the guidelines established by the European Community Council Directive (2010/63/EU). Protocols were approved by the local authority veterinary services and authorized by Italian Ministry of Health (authorizations n. 83/2021-PR and 549/2021-PR).

### Animals strains

*CD1* and *C57BL/6* mice weighting around 25-30 gr were employed for intoxication and infection models, charcoal motility test and faecal pellet analysis, behavioural tests and electrophysiological recording.

Mice were maintained under a 12-hour light/12-hour dark cycle and constant temperature in the animal facility of the Department. Water and food were available ad libitum, with some exceptions (see below). Mice were fed with regular chow.

### Antibodies, reagents and toxins

For Western Blot experiments, cell samples were lysated and loaded in precast polyacrylamide gels (NuPage 4-12%, Bis-Tris Mini protein gel, Invitrogen) immersed in MOPS SDS running buffer (cat. NP0001, Invitrogen novex NuPAGE). Nitrocellulose membranes (cat. 1215471, GVS Life Sciences) were used for protein transfer, and Ponceau S (cat. P7170, Sigma Aldrich) to evaluate protein transfer. Non-fat dried milk powder (A0830.1000, Panreac AppliChem) was used for membrane saturation. The antibodies used were anti-SNAP25 (SMI81) (1:10000, cat. 836303, Biolegend), anti-VAMP2 (1:2000, cat. 104 211, SYSY), anti-cleaved VAMP (1:1000, produced in our laboratory) anti-beta3 tubulin (1:10000, cat. 302302.00, SYSY) anti-mouse IgG HRP-conjugated (1:2000, cat. 401215, Calbiochem), anti-rabbit IgG HRP-conjugated (1:2000, cat. 401393.00, Calbiochem). Membranes were developed using Immobilion Crescendo Western HRP substrate (cat. WBLUR0500, Sigma Aldrich). For immunofluorescence stainings, we used anti-cleaved VAMP (1:1000), anti-cleaved SNAP25 (1:500), both produced in our laboratory, anti-Vesicular Acetylcholine Transporter (VAChT) (1:500, cat. 139 105, primary antibody, Synaptic System), anti-tyrosine hydroxylase (TH) (1:1000, cat. ab76442, primary antibody, Abcam), anti-Substance P (SP) (1:500, cat. PA5-106886, primary antibody, Thermo Scientific), anti-Vasointestinal Peptide (VIP)(1:200, cat. MAB6079-SP, primary antibody, R&D System), anti-WGA fluorescein conjugated (mucin) (1:300, cat. FL-1021, Vector Laboratories), α-bungarotoxin Alexa488-conjugated (1:200, cat. B35451, Life Technologies), anti-guinea pig Alexa555-conjugated (1:200, cat. A-21345, secondary antibody, Thermo Scientific), anti-rabbit Alexa488-conjugated (1:200, cat. A-11090, secondary antibody, Life Technologies), anti-mouse Alexa555-conjugated (1:200, cat. A-21422, secondary antibody, Life Technologies), DAPI (1:5000, cat. D3571, Invitrogen). For cell cultures intoxications, purified BoNT/A and BoNT/B (kindly provided by the laboratory of Professor Shone, CAMR UK) were kept at -80°C and then diluted in cell culture media prior to use. *Clostridial* cultures’ supernatant from *C. botulinum* ATCC 19397 and 27765 (containing BoNT/A and BoNT/B, respectively) were kept at -80°C and diluted in gelatin buffer (0.2% gelatin) prior to use.

The medium used for Cerebellar Granule Neurons cultures is BME (Life Technologies) supplemented with 10% FBS (Euroclone), 25 mM KCl, 2 mM glutamine and 50 μg/ml gentamycin.

For EndoPep-MS assay, Pep-21 for detection of BoNT/A (Ac-RGSNKPKIDAGNQRATR-Nle-LGGR) and Pep-B (H-LSELDDRADALQAGASQFETSAAKLKRKYWWKNLK-OH) were synthesized and purchased from GenScript Biotech (Piscataway, NJ, USA). Antibodies RAZ1 and CR2 (1:1, detection of serotype A) and 2B18.2 and B12.2 (1:1, detection of serotype B) were kindly provided by Dr James D. Marks (University of California, San Francisco, CA, USA). Zeba^TM^ Spin Desalting Columns and sulfo-NHS-biotin were purchased from Thermo Fisher Scientific (Rockford, IL, USA). All reagent for the MALDI Biotyper analysis were acquired from Bruker Daltonics (Bremen, Germany). Magnetic Dynabeads M-280 streptavidin were obtained from Invitrogen-Life technologies (Life Technologies AS, Oslo, Norway).

### Cerebellar granule neurons cultures

5-days-old Sprague-Dawley rats were anesthetized and sacrificed by decapitation. Primary Cerebellar Granule Neurons (CGNs) were obtained as described elsewhere^76^. Briefly, cerebella were collected and then disrupted mechanically using a scalpel. Next, cerebellar tissues were digested using trypsin, and then seeded onto 24-wells culture plates coated with poly-L-lysine (10 μg/ml). Cytosine arabinoside (10 μM) was added to the culture medium 18-24 hours after plating to arrest the growth of non-neuronal cells. Experiments with BoNTs were performed at 6 days in-vitro by intoxication with either purified or supernatant-derived toxins for twelve hours. After the intoxication, CGNs cultures were washed in PBS and lysated for subsequent Western blot analysis.

### Western blotting

After BCA quantification, lysates samples were boiled for 10’. Next, samples were loaded in polyacrylamide gels immersed in diluted MOPS SDS running buffer for protein electrophoretic separation. Proteins were then transferred onto nitrocellulose membranes (pore size 0.45µm) and incubated in Ponceau red solution to ensure that protein transfer has happened properly. After 60’ of saturation, membranes were incubated overnight at 4°C with the proper primary antibodies diluted in BSA solution. The next day, after PBS washes, membranes were incubated for 60’ with proper secondary antibodies and finally developed for samples comparison.

### Mice intoxications

*CD1* mice (or *C57BL/6* mice for infection experiments) weighting 25-30 gr were gavage fed with different volumes of *Clostridial* culture supernatants depending on the dose of BoNT/A or BoNT/B toxins we wanted to obtained. If mice showed signs of botulism, they were sacrificed by cervical dislocation.

### Detection of botulinum toxin in serum samples

Thirty-six hours after intoxication with low doses of BoNT/A or BoNT/B, mice were sacrificed by anaesthetic overdose. Blood samples were collected from the heart into non-heparin-coated tubes, kept at 4°C and centrifuged 4 minutes at 3000g for phase separation. The resulting serum samples were then processed to determine the concentrations of circulating toxin by EndoPep-MS assay^38,39^. Briefly, serum samples (500 µL) were incubated with 20 µL of antibody-coated beads for 1 hour at room temperature on a SB2 fixed speed rotator. Beads were magnetically captured, washed twice with 1 mL PBST, once with 150 µL PBST, and finally with 80 µL ultrapure water. Beads were resuspended in 18 µL reaction buffer (0.1 M DTT, 0.2 mM ZnCl_2_, 1.0 mg/mL BSA, 10 mM HEPES buffer pH 7.3) buffer added with 2 µL of 15 mM specific peptide solution and incubated at 37°C for 4 hours. One microliter of each sample was analyzed using a MALDI Biotyper sirius instrument with 1 µL alpha-cyano-4-hydroxycinnamic acid (HCCA) matrix. The laser frequency was set to 200 Hz, and the final spectrum was an average of 1000 laser shots (10 groups of 100 shots collected in a spiral pattern).

### Compound muscle action potential (CMAP) recording in whisker pad and gastrocnemius

The electrophysiological measurements were performed in 6-8 week old CD1 mice weighting 20-25 gr, anesthetized with a cocktail of xylazine (48 mg/Kg) and zoletil (16 mg/ Kg) via i.p. injections.

Following general anaesthesia, the facial nerve was exposed without damaging the musculature; a small piece of parafilm was put under the nerve, which was kept wet by a drop of physiological solution. A pair of stimulating needle electrodes (Grass, USA) were advanced until they gently touched the exposed nerve using a mechanical micromanipulator (MM33, FST, Germany). A pair of needle electrodes for electromyography (Grass, USA) were used for electromyographic recording of whisker pad muscle or hind limb muscle fibres’ activity. In whisker pad’s analysis, the recording needle electrode was inserted halfway in the whisker pad while the indifferent needle electrode was inserted under the skin of the nose. In hind limbs’ analysis, the recording needle electrode was inserted halfway in the gastrocnemius, while the indifferent needle electrode was inserted under the skin of the paw. The analysis was performed 72 or 168 hours after intoxication with BoNT/A or BoNT/B. Compound muscle action potentials (CMAPs) were recorded upon supramaximal stimulation of the facial or sciatic nerve (25V, 4ms, rate of stimulation of 0.5Hz) using a stimulator (S88, Grass, USA) via a stimulus isolation unit (SIU5, Grass, USA) in a capacitive coupling mode. Recorded signals were amplified by an extracellular amplifier (P6 Grass, USA), digitized using a digital A/C interface (National Instruments, USA) and then fed to a PC for both online visualization and offline analysis using appropriate software (WinEDR, Strathclyde University; pClamp, Axon, USA). Stored data were analyzed offline using pClamp software (Axon, USA).

### Rotarod test

For rotarod test, mice were trained for five days before the intoxication with sBoNT. The protocol was decided based on the literature^77–79^ and on previous experience of our laboratory. Mice were placed on the apparatus in a quiet, dark room. The apparatus was set up to increase rotation speed (5 to 30rpm) over a time of 240 seconds. If the mice felt from the rotating tube or cling on it and rotate for more than two rotations, the test was considered failed and data were recorded. Data were then analyzed as amount of time spent walking and reported as percentage of test accomplished.

### Ventilation recording

Following general anesthesia, animals were left 10 minutes in their cages to fully relax. For each mouse, recordings were performed before (t0) and 72 or 168 hours after intoxication with sBoNT/A or sBoNT/B. A 20 ga x 38 mm plastic feeding tube (Instech Laboratories, Inc.) connected to a pressure sensor (Honeywell, 142PC01D) was carefully introduced in the mice oral cavity and placed in the lower third of the esophagus at the level of mediastinum. The animals laid on their left side on a pre-warmed surface to record their mediastinal pressure variations, which were used to infer animal ventilation. Traces were recorded, amplified and digitized with WinEDR V3.4.6 software (Strathclyde University, Scotland). Stored data were analyzed using Clampfit software (Axon, USA). For each animal, 120 epochs were recorded and 20 epochs were analyzed at each time point. Inferred Ventilation Index (IVI) was then calculated as the product of the mean area of the peaks (mV x ms) and the number of peaks within 20 sec, and represented in the graphs as the % of recovery of the animals compared to their own initial record (t0). We indirectly detected the volume of air flux by measuring esophageal pressure variations, which reflect intrapleural pressure variations^80^ and respiratory muscles wellness.

### Charcoal motility test

CD1 mice were kept under starvation for 12 hours and then gavage-fed with a bolus of charcoal (10% w/v) and Arabic gum (5% w/v) dissolved in water. Mice were caged individually, and the time needed for the expulsion of first black, faecal pellet was assessed. The collection of faecal pellets continued until 200 minutes from the start of the experiment. The same mice underwent the measurement before (vehicle) and after treatment with sBoNT/A or sBoNT/B. High-dose treated animals failed the test (>200 minutes) and were excluded from the analysis.

### Faecal pellet humidity measurment

After mice treatment either with sBoNTs or vehicle, a single faecal pellet was sampled and weighted at different timepoints. After overnight incubation at 60°C^81,82^, faecal samples were weighted again and the relative content of faecal moisture was determined.

### Infection models

Sixty hours after the oral administration of the vehicle, sBoNT/A, or sBoNT/B, mice were treated with a broad-spectrum antibiotic solution consisting of 50 mg/kg vancomycin, 100 mg/kg neomycin, 100 mg/kg metronidazole, and 100 mg/kg ampicillin by oral gavage. Such treatment was already performed in our laboratories without showing changes in animals in terms of intestinal motility (no appearance of diarrhoea or constipation, no signs of blood in stools), clinical pain signs (hunched posture, ruffled fur) and eating habits (equal weight of food consumed). The treatment with the antibiotic mixture was necessary to reduce the normal intestinal microbiota of the animal and promote the adhesion of the microbial load of *Salmonella enterica serovar typhimurium* (*S. typhimurium*, 1 × 10^8^ CFU in 100 μl) and *Shigella flexneri serovar 5a/M90T* (*S. flexneri*, 1 × 10^7^ CFU in 100 μl) that were administered 12 hours later by oral gavage.

Five days later, the animals were sacrificed by cervical dislocation and Peyer’s patches, spleens and livers were collected, homogenized, diluted and plated on SS selective agar. The plates were then incubated at 37°C for up to 36 hours, and CFUs were counted. Small intestines were collected for following immunofluorescence analysis.

### Samples preparation for immunofluorescence stainings

Mice were sacrificed after the electrophysiological and infection experiments aforedescribed, according to European guidelines for animal care, and different tissues were collected. Whisker pad muscles were fixed for 30’ in PBS + 4% Paraformaldehyde (PFA) at room temperature (RT), dried and then included in OCT and frozen. Gastrocnemius and diaphragm muscles were instead dissected and fixed in PBS + 4% PFA (30’ at RT), and then used for immunofluorescence (IF). Small intestines were collected, fixed 1 hour in PBS added with 4% Paraformaldehyde (PFA) at room temperature (RT) and then used for whole mount immunofluorescence. Otherwise, they were rolled, fixed overnight (ON) in PBS added with 4% PFA and 15% sucrose (weight/volume), transferred in PBS + 30% sucrose for 48 hours and then included in OCT and frozen. Whisker pad muscles, Peyer’s Patches and small intestines samples included in OCT were then cryosliced at 15 µm and then used for IF stainings.

### Immunofluorescence stainings

All tissues were quickly washed in PBS for three times, five minutes each, and then incubated in PBS added with 50mM NH_4_Cl for quenching. Then, tissues were incubated in PBS added with 0.5% Triton X100 and 3% BSA for permeabilization and saturation. Next, they incubated ON or for two days (for whole mount stainings) at 4°C with different primary antibodies (diluted in the same solution of the previous passage). The next day, tissues were washed three times in PBS, and then incubated with the required secondary antibodies (diluted in PBS) for 2 hours at RT. Finally, the tissues were washed twice with PBS, incubated for 5’ at RT with DAPI, washed another time in PBS and mounted on proper glasses using DAKO mounting solution.

For cell cultures stainings, cells were fixed 10’ in 4% PFA (wt/vol) and washed twice in PBS. Cells were then quenched 10’ in 50 nM NH_4_Cl, and permeabilized for 3 min at RT with 0.3% Triton X-100 in PBS. After saturation with 3% Goat serum in PBS for 1 h, samples were incubated with primary antibody diluted in the same solution ON. After washings and incubation with corresponding Alexa-conjugated secondary antibodies for 1 hour at RT, coverslips were mounted using Fluorescent Mounting Medium.

### Microscopy and image analysis

Images were collected with a confocal microscope (ZEISS LSM900 airyscan2) equipped with ECPlan-Neofluar (20X/0.5 air, 40X/0.45 oil) or Plan-Apochromat (63X/1.46 oil, 100X/1.4 oil) objectives for z/stacks acquisitions. Images were then processed using high resolution airyscan processing, and analyzed using ZEN 3.4 (blue edition) or ImageJ programs. To ensure the absence of background noise and non-specific staining puncta in SNARE cleavage detection, ImageJ “remove noise outliars” tool was used, using stainings from vehicle-treated enteric samples as cut-off. Mucin average signal intensity was measured using ImageJ “measure” tool. For each intestine sample, the signal intensity was measured in three different areas, and ten villi/area were analyzed. Each dot represents the average signal intensity of an entire area.

### Statistical analysis

Sample sizes were determined by analysis based on data collected by our laboratory in published studies. For ventilation recording and charcoal motility test, we applied the techniques to the same mice before and after the treatment with sBoNTs. N = 6 mice/group for low-sBoNT/A and low-sBoNT/B and n = 3 for high-sBoNT/A and high-sBoNT/B were used in electrophysiological tests, charcoal motility test, faecal pellet moisture measurement, mice weight and food consumption measurements. For foodborne infection models, n = 5 mice/group were used. We ensured a blind conduct of experiments. GraphPad Prism software was used for statistical analyses. Statistical significance was evaluated using unpaired Mann-Whitney test, by One-way analysis of variance (ANOVA) with Tukey’s post-hoc test or with by Wilcoxon signed-rank test (expected value = 0 for ventilation test, expected value = 100 for rotarod test). Data were presented as median values with interquartile range, and considered statistically different when *p < 0.05, ** p < 0.01, *** p < 0.001, **** p < 0.0001.

## Acknowledgements

This research was supported by grants from the Ministry of Defence “RIPANE project” (to CM) and by the University of Padua “Progetto DOR 205071” (to OR).

## Author contribution

F.F., P.B., G.B., I.C., I.D. performed the experiments and analysed the data. C.M. and O.R. conceived the study and wrote the manuscript. A.M., L.B., F.L., M.L.B. provide reagents and assisted in interpreting and discussing the results. All the authors reviewed the manuscript.

## Conflict of interest

The authors declare no conflicts of interest.

## Supplementary Figure Legends

**Figure S1.**
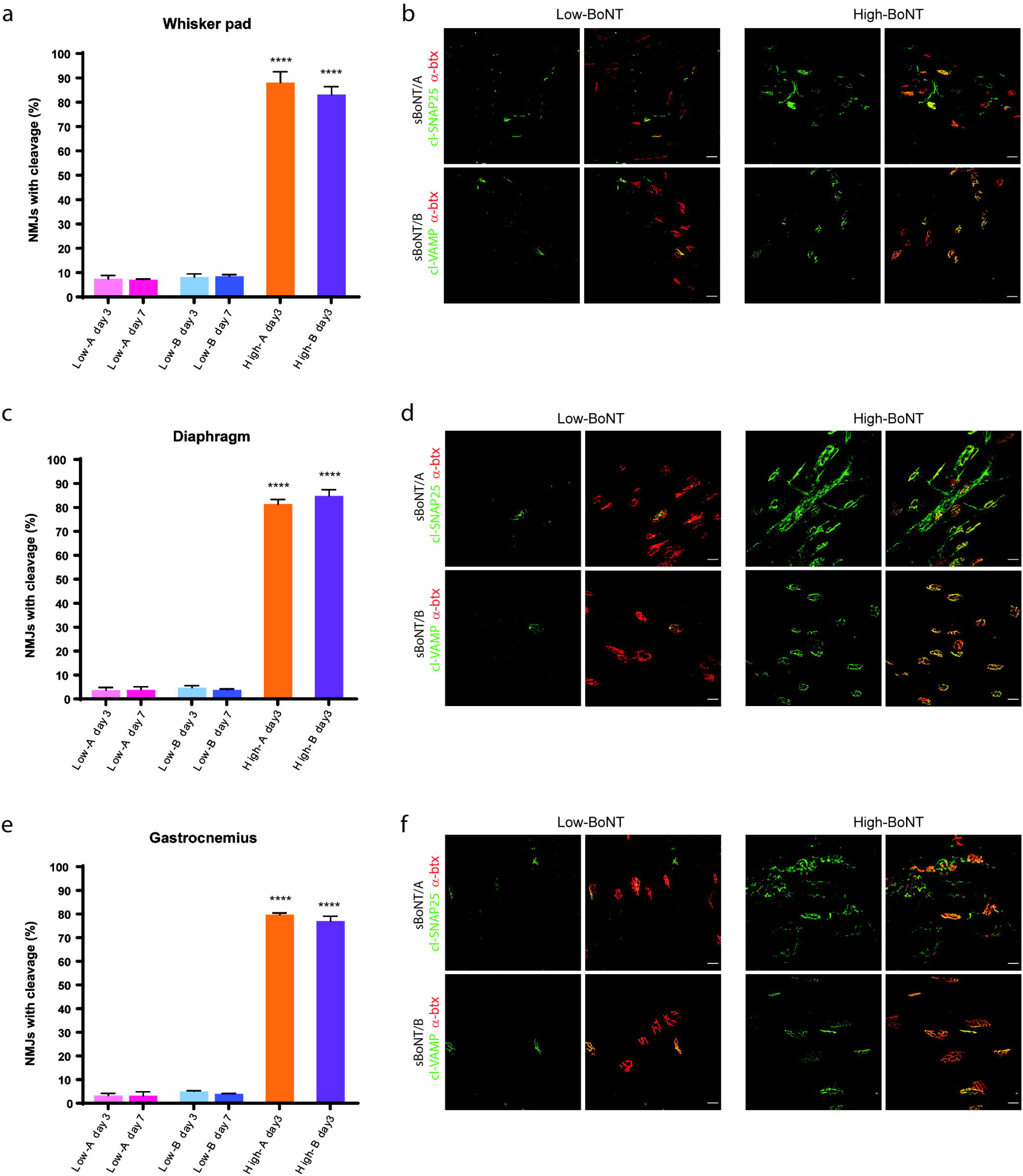
Analysis of SNAP25 and VAMP cleavage in neuromuscular junctions of mice intoxicated with high or low doses of sBoNT. **a.** Quantification of the number of neuromuscular junctions (NMJs) in the whisker pad with staining for cl-SNAP25 (sBoNT/A) and cl-VAMP (sBoNT/B), reported as a percentage of the total NMJs counted. Data from mice treated with low doses of sBoNT/A are shown in pink or fuchsia (count at three and seven days post-intoxication, respectively), with low doses of sBoNT/B shown in light blue or blue, while data from intoxication with high doses of sBoNT/A or sBoNT/B at three days post-intoxication are shown in orange and purple, respectively. **b.** Representative staining of cl-SNAP25 and cl-VAMP in NMJs of whisker pad muscles from mice intoxicated with BoNT. Staining for cl-SNAP25 (upper panels) and cl-VAMP (lower panels) is shown in green, while staining for nicotinic receptors at the NMJ (α-btx) is shown in red. The left panels show staining in muscles from mice treated with low doses of toxin, while the right panels show staining from mice treated with high doses of toxin. **c.** Quantification of the number of NMJs in the diaphragm with staining for cl-SNAP25 (sBoNT/A) and cl-VAMP (sBoNT/B), reported as a percentage of the total NMJs counted. Data in pink or fuchsia refer to mice intoxicated with low doses of sBoNT/A (count at three and seven days post-intoxication, respectively), those shown in light blue or blue refer to low doses of sBoNT/B, while data from intoxication with high doses of sBoNT/A or sBoNT/B at three days post-intoxication are shown in orange and purple, respectively. **d.** Representative staining of cl-SNAP25 and cl-VAMP in NMJs of diaphragm muscles from mice intoxicated with BoNT. Staining of muscles from mice treated with low doses of sBoNT is shown in the left panels, while those treated with high doses are shown in the right panels. Staining for cl-SNAP25 (upper panels) and cl-VAMP (lower panels) is shown in green, while staining for nicotinic receptors at the NMJ (α-btx) is shown in red. **e.** Quantification of the number of NMJs in the gastrocnemius with staining for cl-SNAP25 (sBoNT/A) and cl-VAMP (sBoNT/B). Data from mice treated with low doses of sBoNT/A are shown in pink or fuchsia (count at three and seven days post-intoxication, respectively), with low doses of sBoNT/B shown in light blue or blue, while data from intoxication with high doses of sBoNT/A or sBoNT/B at three days post-intoxication are shown in orange and purple, respectively. The data are reported as the percentage of NMJs with cleavage staining relative to the total NMJs counted. **f.** Representative staining of cl-SNAP25 and cl-VAMP in NMJs of gastrocnemius muscles from mice intoxicated with BoNT. Cleavage staining in mice treated with sBoNT/A (cl-SNAP25, upper panels) and sBoNT/B (cl-VAMP, lower panels) is shown in green, while staining for nicotinic receptors at the NMJ (α-btx) is shown in red. The left panels show staining in muscles from mice treated with low doses of toxin, while the right panels show staining from mice treated with high doses of toxin. Images acquired at 20X, scale bar: 25μm. Statistical analysis was performed using one-way analysis of variance (ANOVA) with Tukey’s post-hoc test and multiple comparisons. ****p < 0.0001.

**Figure S2.**
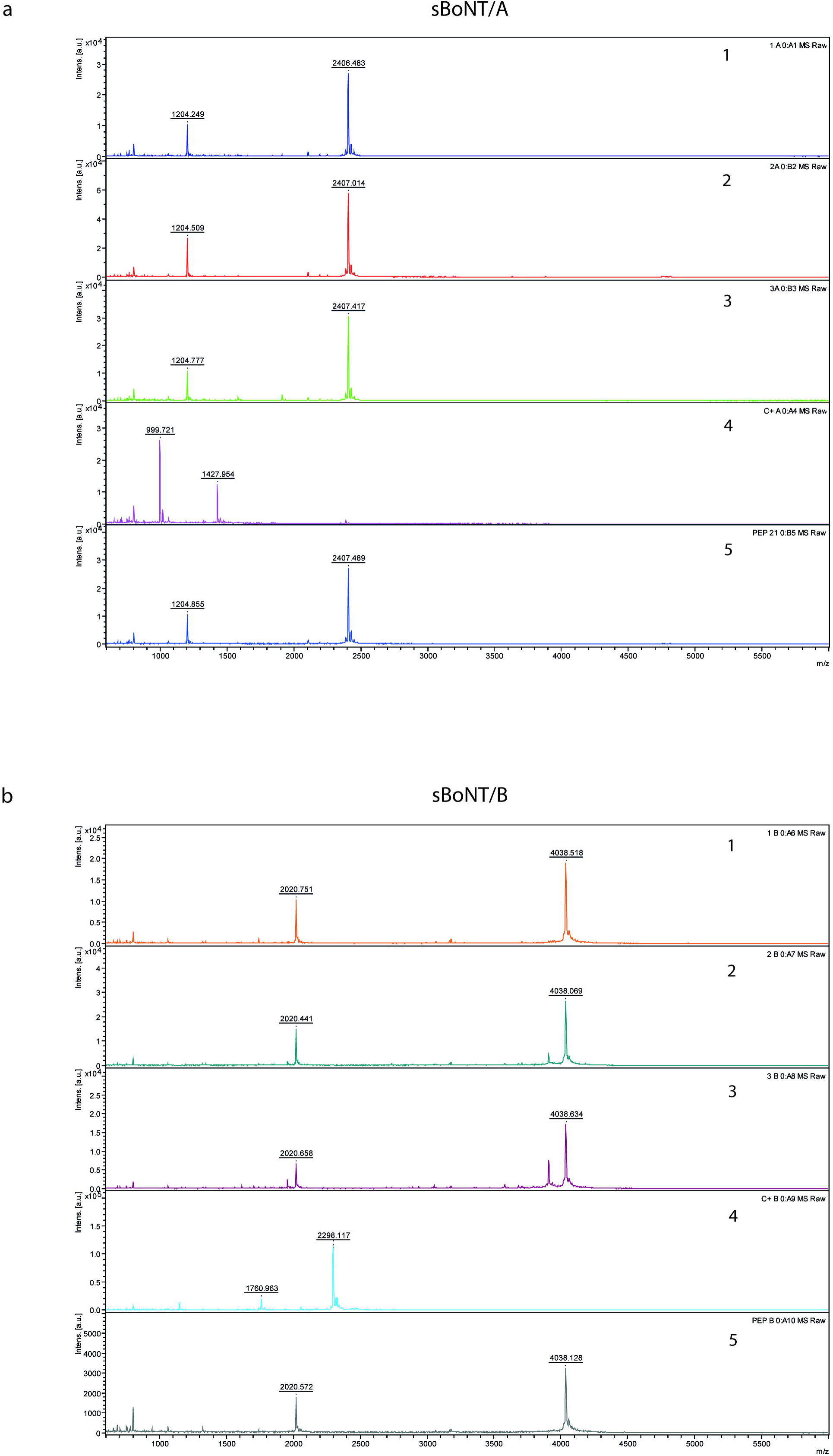
EndoPep-MS analysis of the serum from BoNT-intoxicated mice. **A.** BoNT/A detection using EndoPep-MS on sera collected from mice. (1), (2) and (3) spectra obtained with sera collected from mice orally intoxicated with the low dose of sBoNT/A: only the intact, mono- and bi-protonated peptide was detected (1204.855 and 2407.489 *m/z* respectively). No evidence of BoNT/A was found. (4) Spectrum from the reaction of sera spiked with both BoNT/A (positive control) and pep-21 showed cleavage products of pep-21 at m/z 999.721 and 1427.954, respectively, confirming the presence of BoNT/A in the sample. (5) Negative sera spiked only with the pep-21 showed spectra with a single charged ion at m/z 2407.489 and a double charged ion at m/z 1208.555. **B.** BoNT/B detection using EndoPep-MS on sera collected from mice. (1), (2) and (3) spectra obtained with sera collected from mice intoxicated with the low dose of sBoNT/B: only the intact, mono- and bi-protonated peptide was detected (4038.518 and 2020.751 *m/z* respectively). No evidence of BoNT/B was found. (4) Spectrum from the reaction of sera spiked with both BoNT/B (positive control) and pep-B showed cleavage products of pep-B at *m/z* 1760.963 and 2298.117, respectively, confirming the presence of BoNT/B in the sample. (5) Negative sera spiked only with the pep-21 showed spectra with a singly charged ion at *m/z* 4038.518 and a doubly charged ion at *m/z* 2020.751.

**Figure S3.**
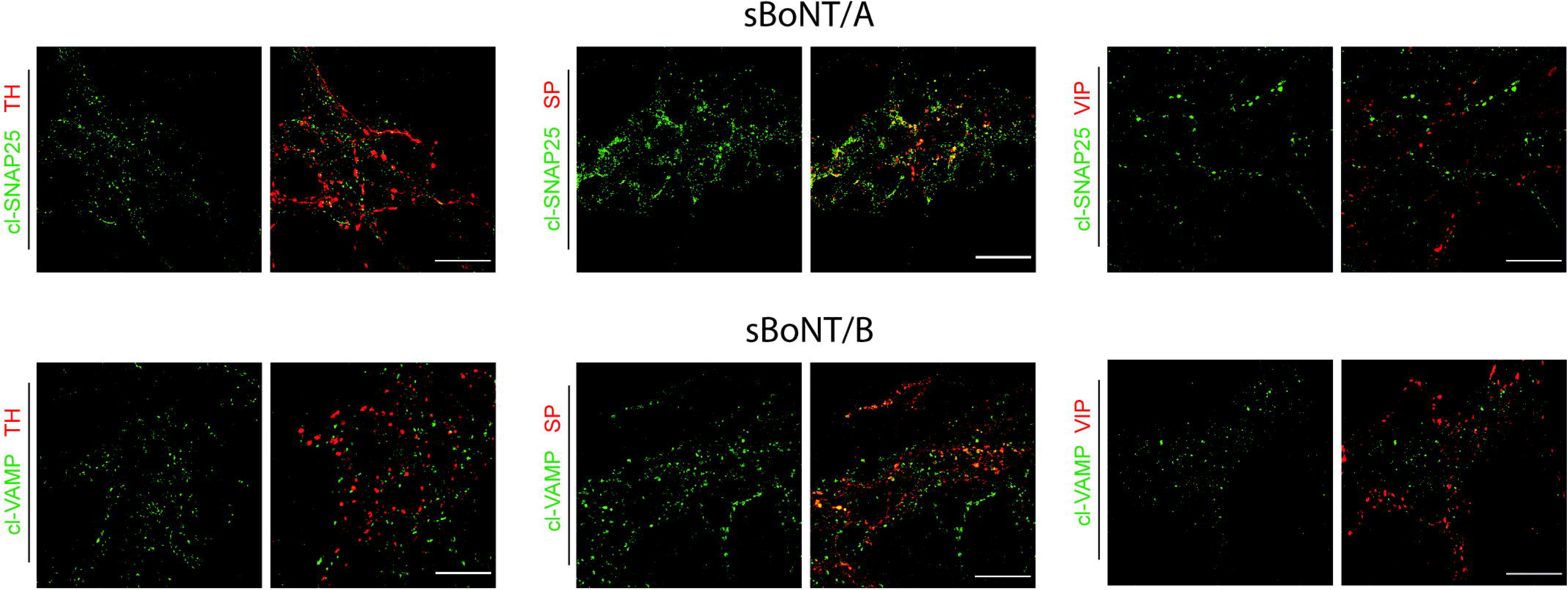
Analysis of the proteolytic activity of BoNT in non-cholinergic enteric neurons. Staining of cleavage of SNAP25 (upper panels) and VAMP (lower panels) in the myenteric plexus of mice intoxicated with low doses of sBoNT/A and /B, respectively. The left images show co-staining with the marker for tyrosine hydroxylase (TH), the central images show co-staining with the marker for Substance P (SP), while the right images show co-staining for Vaso-Intestinal Peptide (VIP), all three in red. Images were acquired at 63X, scale bars: 25μm.

## Notes

### Competing Interest Statement

The authors have declared no competing interest.

